# Reconstitution reveals friction-driven membrane scission by the human ESCRT-III proteins CHMP1B and IST1

**DOI:** 10.1101/2022.02.03.479062

**Authors:** A. King Cada, Mark R. Pavlin, Juan P. Castillo, Alexander B. Tong, Kevin P. Larsen, Xuefeng Ren, Adam Yokom, Feng-Ching Tsai, Jamie Shiah, Patricia M. Bassereau, Carlos J. Bustamante, James H. Hurley

## Abstract

The endosomal sorting complexes required for transport (ESCRT) system is an ancient and ubiquitous membrane scission machinery that catalyzes the budding and scission of membranes. ESCRT-mediated scission events, exemplified by those involved in the budding of HIV-1, are usually directed away from the cytosol (‘reverse-topology’), but they can also be directed towards the cytosol (‘normal-topology’). Of the ESCRT complexes 0-III, ESCRT-III is most directly implicated in membrane severing. Various subunits of ESCRT-III recruit the AAA^+^ ATPase VPS4, which is essential for ESCRT disassembly and reverse topology membrane scission. The ESCRT-III subunits CHMP1B and IST1 can coat and constrict positively curved membrane tubes, suggesting that these subunits could catalyze normal topology membrane severing, perhaps in conjunction with a AAA^+^ ATPase. CHMP1B and IST1 bind and recruit the microtubule-severing AAA^+^ ATPase spastin, a close relative of VPS4, suggesting that spastin could have a VPS4-like role in normal topology membrane scission. In order to determine whether CHMP1B and IST1 are capable of membrane severing on their own or in concert with VPS4 or spastin, we sought to reconstitute the process *in vitro* using membrane nanotubes pulled from giant unilamellar vesicles (GUVs) using an optical trap. CHMP1B and IST1 copolymerize on membrane nanotubes, forming stable scaffolds that constrict the tubes, but do not, on their own, lead to scission. However, CHMP1B-IST1-scaffolded tubes were severed when an additional extensional force was applied, consistent with a friction-driven scission mechanism. Spastin colocalized with CHMP1B enriched sites but did not disassemble the CHMP1B-IST1 coat from the membrane. VPS4 resolubilized CHMP1B and IST1 but did not lead to scission. These data show that the CHMP1B and IST1 tubular coat contributes to membrane scission. Constriction alone is insufficient for scission. However, the dynamical extension of the coated tube does lead to scission. Finally, we find that in the normal topology setting analyzed here, scission is independent of VPS4 and spastin. These observations show that the CHMP1B-IST1 ESCRT-III combination is capable of severing membranes by a friction-driven mechanism.

## Introduction

The Endosomal Sorting Complexes Required for Transport (ESCRT) proteins are an ancient and conserved membrane remodeling machinery, present in two of the three domains of life, the Archaea and Eukaryota. In humans, the ESCRTs are involved in myriad of cell biological processes (1, 2) ranging from multivesicular body biogenesis (3), cytokinetic abscission (4), membrane repair (5-8), and exosome (9) and HIV-1 release (10, 11). The underlying commonality of most of these processes is that they are topologically equivalent, with scission occurring on the cytosolic and inner surface of a narrow membrane neck (‘reverse topology’). The ESCRTs consist of ALIX, ESCRT-0, ESCRT-I, ESCRT-II, ESCRT-III, and VPS4 (12, 13). The ESCRT-III proteins (14) are the ones most directly involved in catalyzing membrane scission (15, 16). These ESCRTs are first recruited to the neck then the AAA^+^ ATPase VPS4 (17) is finally recruited to ESCRT-III enriched sites prior to scission (18). VPS4 forms a hexamer (19, 20) that interacts with ESCRT-III through its N-terminal microtubule-interacting and trafficking (MIT) domain binding to the exposed C-terminal MIT domain interacting motifs (MIM) domains found in some ESCRT-III proteins (21, 22). ESCRT-III together with VPS4 constitutes the minimal module to drive scission of vesicles that bud away from the cytosol (16). While ESCRTs are best known for reverse-topology membrane scission, a subset of ESCRTs, CHMP1B and IST1, can also coat the outer surface of membrane tubes, leading to a dramatic constriction in the tube (23, 24). This process is implicated in tubular endosomal traffic from the ER to lysosomes (25-27) and lipid droplets to peroxisomes (28), and the release of newly formed peroxisomes from the ER (29). These observations suggested that CHMP1B and IST1 could carry out normal topology scission; however, direct observation of this type of scission has not been reported.

Twelve different ESCRT-III proteins are found in humans, which can combine in various compositions that nucleate and grow on membranes of various curvatures (23, 30, 31). ESCRT-III proteins are monomeric and are highly basic, and share similar secondary core structures containing five helices (32). These proteins are in an auto-inhibited closed conformation in solution (33, 34). Activation can be triggered upon binding to membranes or upstream activators, or artificially through truncation of their C-terminal elements (33, 34). Upon activation, ESCRT-III proteins polymerize into spirals (35-37), and helical tubes (24, 38). Incubation of CHMP1B with liposomes lead to formation of protein coated tubules *in vitro* as shown using cryo-electron microscopy (cryo-EM) (23). This positively curved coat was initially unexpected in the ESCRT field, but subsequently has been observed more generally with combinations of CHMP2, 3, and 4 (39, 40).

In reverse-topology scission, the ATPase activity of VPS4 is essential for the remodeling of ESCRT-III assemblies that lead to membrane constriction and scission (16). Polymerization of CHMP4 is considered a major driver of scission, however, this is held in check by capping with CHMP2 (41). VPS4 can solubilize CHMP2 subunits, allowing CHMP4 growth to progress, leading to scission (42). VPS4 binds to most CHMPs, including IST1 and CHMP1; however, CHMP2 is not known to have a role in normal topology scission, and CHMP1 and IST1 are not known to engage in capping. Therefore, it is not clear if VPS4-driven decapping mechanism established in reverse topology membrane scission has a role in normal topology scission.

Spastin belongs to the same meiotic clade of AAA^+^ ATPases as VPS4, and is best known as a microtubule severing enzyme (43, 44). Mutations of the spastin (SPG4, SPAST) gene are the main causes in patients suffering from hereditary spastic paraplegia (45). The spastin linkage to CHMP1B and IST1 is involved in the scission of recycling endosomal tubules (25). Disruption of any of these interactions increases endosomal tubulation, mistrafficking of cargoes, and dysregulation of proper lysosomal functions (25-27). It has remained unclear if spastin can substitute for the possible scission or disassembly functions of VPS4 with respect to CHMP1B and IST1 containing membrane tubes, in addition to its canonical microtubule severing activity.

Recent cryo-EM reconstructions on synthetic liposomes showed that the polymerization of IST1 on the exterior of CHMP1B-coated tubes leads to a remodeling of the CHMP1B coat and tightly constricts the membrane, but does not lead to membrane scission (24). Here, we assayed ESCRT membrane scission *in vitro* by combining optical tweezers and fluorescence microscopy to visualize membrane nanotubes pulled from giant unilamellar vesicles (GUVs) (46, 47) and characterize the effect in addition of various ATPase on ESCRT dynamics. This assay allows for the formation of a positively-curved membrane that mimics the membrane tubules where CHMP1B and IST1 bind. We were able to reconstitute the scission reaction and delineate the roles of CHMP1B, IST1, VPS4, and spastin in normal topology scission.

## Results

### Binding and constriction of CHMP1B and IST1 does not cause scission

We purified full-length CHMP1B and a truncated IST1 construct that spans residues 1-189 (IST1^NTD^), which was previously shown to form tubular coats with enhanced helical order (24). CHMP1B and IST1 were incubated together with liposomes and visualized by cryo-EM, verifying that these are the minimal constructs to form protein coated tubules (Fig. 1a). In order to probe the consequences of ESCRT-III binding on the membrane in real time, we employed a membrane tube pulling assay. We formed membrane nanotubes between aspiration pipet-immobilized GUVs (60% egg phosphatidylcholine and 40% dioleoylphosphatidyserine with trace dioleoylphosphatidylethanolamine-Atto488 and distearoylphosphatidylethanolamin-PEG (2000)-biotin) and a streptavidin-coated bead by briefly putting them in contact and then pulling them apart. After pulling a tube, a second pipette filled with 5 μM ESCRT-III proteins LD555-CHMP1B and unlabeled IST1^NTD^ was lowered and its contents dispensed in proximity (20 – 30 μm) to the membrane tube. We monitored the reaction via fluorescence using confocal microscopy (Fig. 1b). Consistent with previous reports, LD555-CHMP1B immediately bound to membranes containing high negative charge density lipids (40 mol% PS) (48) and constricted the membrane nanotubes (Fig. 1c). We observed membranes constricted to 25 ± 6 nm radii (mean ± SD) (Fig. 1d), as measured by lipid fluorescence (Extended Data Fig.1). Although ESCRT-III binding alone constricted membrane tubes in this system, we did not observe scission of ESCRT-III bound nanotubes (*n* = 13) under these conditions.

**Fig.1:**
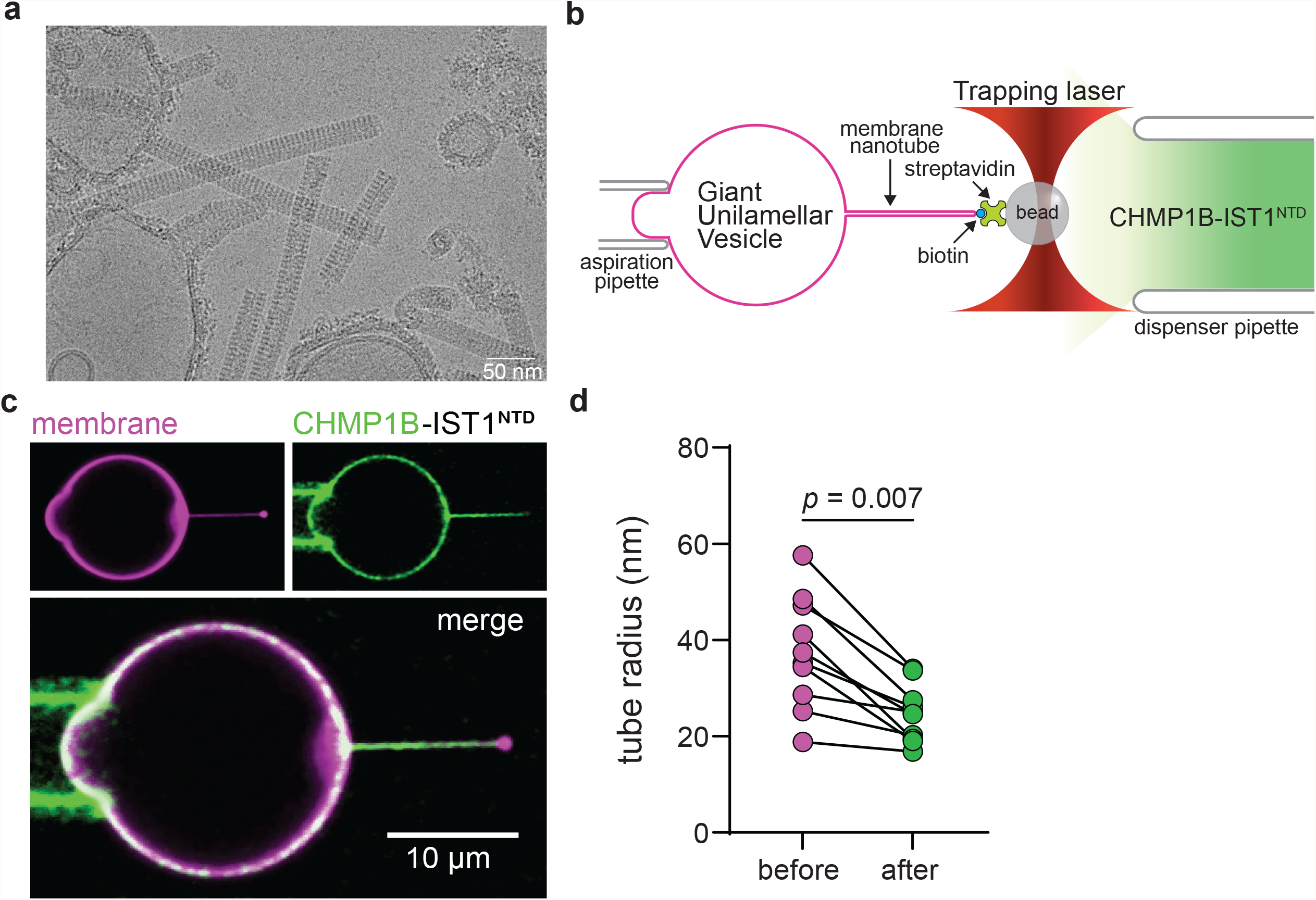
ESCRT-III subunits CHMP1B and IST1^NTD^ bind and constrict membranes. **a**, Representative cryo-EM micrograph of ESCRT-III tubulating the membrane **b**, Fluorescently-labeled giant unilamellar vesicles (GUV) containing 59.4 mol% ePC, 40 mol% DOPS, 0.5 mol% DOPE-ATTO488, and 0.1 mol% DSPE-PEG-2000-biotin were held in place by suction using an aspiration pipette. Membrane nanotubes were formed between the immobilized GUV and a streptavidin-coated bead held by an optical trap after briefly putting them into contact and subsequently retracting them apart. **c**, Representative confocal images showing LD555-CHMP1B (green) and IST1^NTD^ on the tube after addition of 5μm proteins on the membrane labeled with 0.5 mol% DOPE-ATTO488 (magenta). **d**, Membrane tube diameter decreases upon binding of ESCRT-III proteins.

### CHMP1B-IST1 severs membranes upon dynamical tube extension

Since CHMP1B and IST1 can tubulate and scaffold the membrane, we asked whether these polymers are rigid structures that could serve as lipid diffusion barriers and potentially promote friction-driven scission (FDS) (49). Fluorescence recovery after photobleaching (FRAP) was used to measure the dynamics of LD555-CHMP1B and IST1^NTD^ on GUVs. We initially incubated the GUVs with 500 nM LD555-CHMP1B and IST1^NTD^ followed by dilution to remove unbound proteins on the GUV. After incubation, GUVs show homogeneous CHMP1B coverage based on the fluorescence. After photobleaching a region on the GUV (Fig.2a) we observed that fluorescence intensity did not recover after 2 minutes, indicating that the bound CHMP1B is immobile (Fig. 2b). We similarly performed FRAP experiments on the membrane, revealing that CHMP1B and IST1^NTD^ can slow lipid diffusion (Fig. 2c).

**Fig.2:**
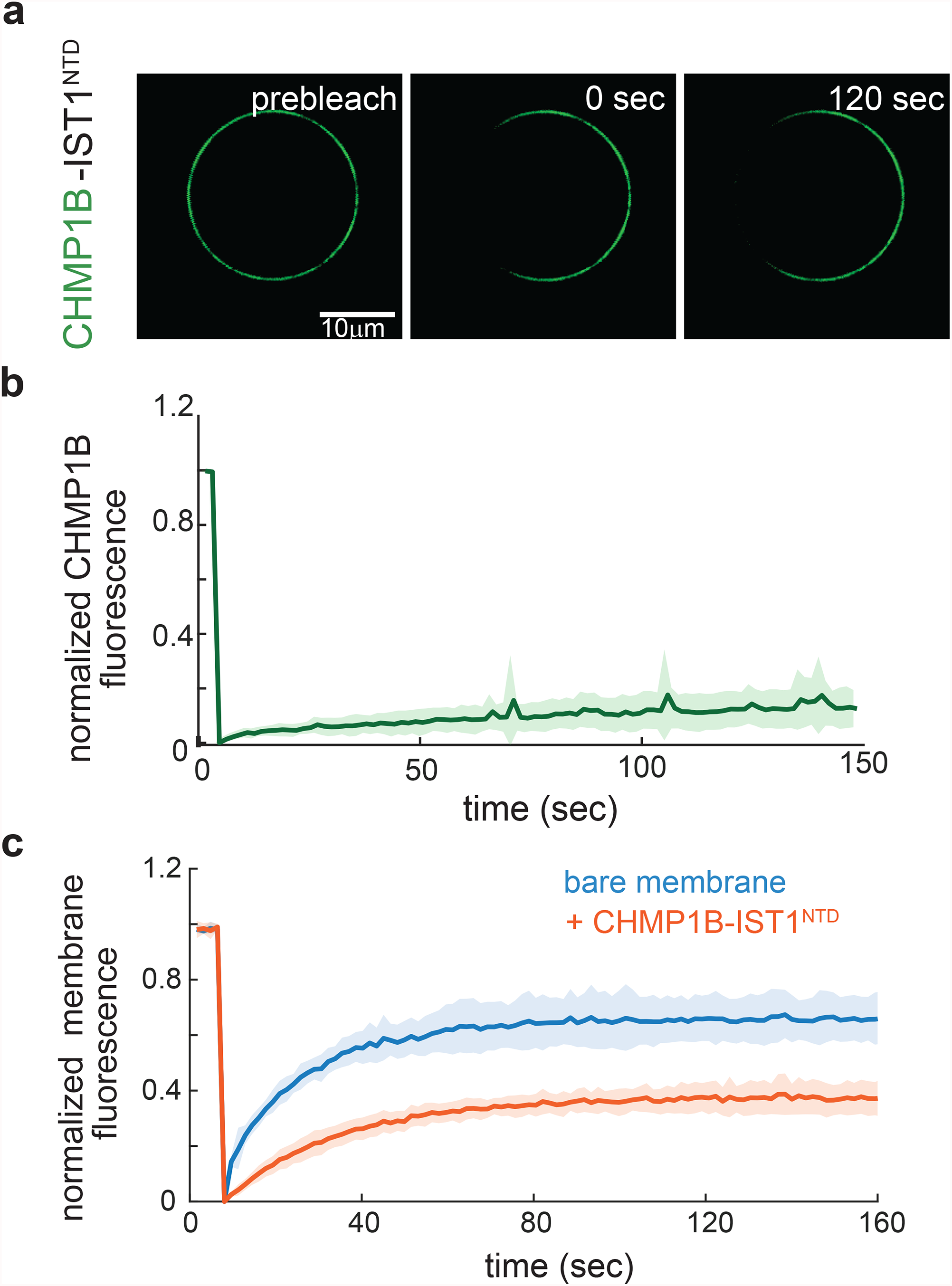
CHMP1B and IST1^NTD^ form rigid structures on the membrane and acts as a lipid diffusion barrier. **a**, Representative image of LD555 labeled CHMP1B before and after performing fluorescent recovery after photobleaching (FRAP) to measure protein mobility on the membrane. 500 nM LD555-CHMP1B and IST1^NTD^ were pre-adsorbed onto the GUV and diluted 5X to avoid recovery from soluble protein on the GUV from the external solution. **b**, Recovery curve of LD555-CHMP1B showing that ESCRT-III subunits are immobile once bound on the membrane. Results are means ± standard deviation from 6 FRAP experiments. **c**, FRAP recovery curve of DOPE-ATTO647 after photobleaching on the GUV shows slow diffusion of lipids when proteins are bound (orange curve) compared to negative control measuring recovery of GUVs in the absence of proteins (blue curve). Results are means ± standard deviation from 6 FRAP experiments.

We next asked whether formation of a rigid ESCRT-III coat can promote scission driven by friction with the tube membrane. To test this, we generated membrane nanotubes from GUVs using optical tweezers as above, and then subjected them to additional back and forth movement along the tube axis. As a control, we did not observe spontaneous scission of bare membranes upon axial motion (Fig. 3a and Supplementary Video 1). After CHMP1B and IST1 binding had been established on the membrane tube, we pushed and pulled the tubes axially as before, and observed scission as evidenced by loss of membrane fluorescence connecting the GUV and the bead. Formation of a rigid protein scaffold that can act as a lipid diffusion barrier on the tube was evident when we pushed the tube closer to the bead (Fig. 2b). Typically, two cycles of pushing and pulling were carried out over a distance of 30 μm at a speed of 3 μm s^-1^. The presence of CHMP1B fluorescence seen at the end of the membrane tube suggests scission occurred at the edge of the protein scaffold (Fig. 2c,d and Supplementary Video 2). Therefore, membrane scission by ESCRT-III proteins can be promoted by applying an additional mechanical force through tube extension in the presence of a rigid protein scaffold.

**Fig.3:**
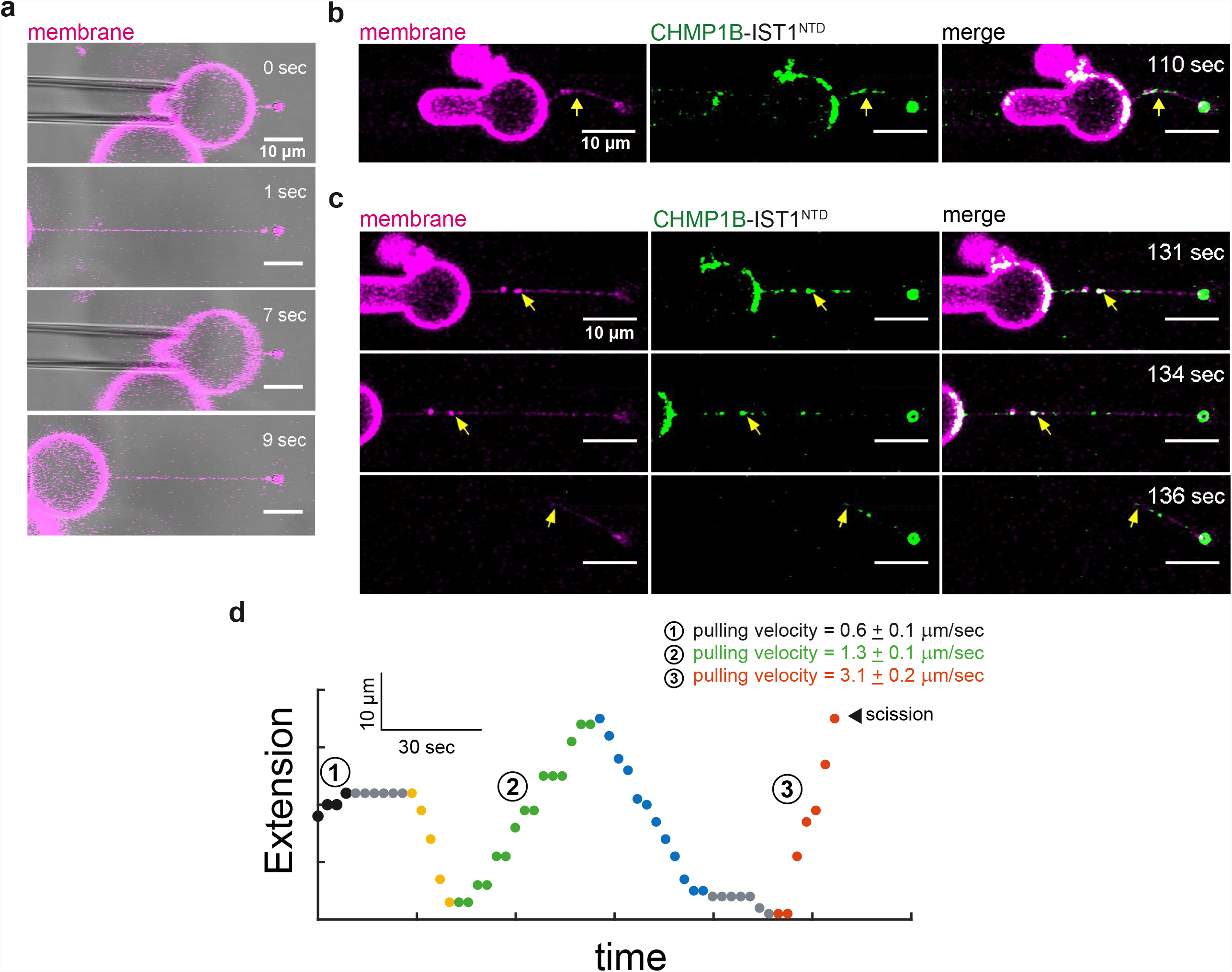
External pulling on ESCRT-III scaffolded tubes promotes scission. **a**, Bare membranes pulled at > 25 μm.s^-1^ at 0.2 pN.nm^-1^ does not break even after repeatedly being brought back-and-forth. **b**, Membrane tube resisted retraction when pushed at 2 μm.s^-1^ after LD555-CHMP1B enrichment. 5 μM LD555-CHMP1B and 5 μM IST1^NTD^ were dispensed using a micropipette in proximity to the region of interest. **c**, Snapshots of membrane tube pulled at 3 μm.s^-1^ on membrane bound LD555-CHMP1B-IST1^NTD^ protein scaffolds induces scission. Yellow arrow highlights the point of scission. **d**, Representative tube pulling trajectories of CHMP1B-IST1^NTD^ protein scaffolds on the tube. Pulling velocities below 3 μm.s-1 does not lead to scission (black and green circles) while pushing tubes at ∼2 μm.s-1 show tube bending in (**b**). Results are means ± standard deviation from 5 experiments Scale bars are 10μm.

### Spastin does not disassemble CHMP1B and IST1 from the membrane

In reverse-topology ESCRT processes, the AAA^+^ ATPase VPS4 is critical in the disassembly and remodeling of ESCRTs (50). VPS4 is thought to drive the constriction and eventual scission (16) of membranes by continuously removing the CHMP2 cap (42). Spastin belongs to the meiotic clade of AAA^+^ ATPase together with VPS4, and is recruited by CHMP1B and IST1 in the endosomal recycling pathway (25, 27). Like VPS4, spastin contains a microtubule interacting and trafficking (MIT) domain that interacts with the MIT –interacting motif (MIM) domain found on the C-terminus of CHMP1B (51) and in full-length IST1. Spastin predominantly exists as two isoforms in mammalian cells and acts primarily on microtubules (43, 52). Full-length spastin (M1-spastin) contains an N-terminal hydrophobic domain which tethers it to the endoplasmic reticulum (53). In contrast, the most abundant isoform, M87-spastin, is cytosolic and can be recruited to microtubules and early endosomes (25). Therefore, we purified M87-spastin and tested the stimulation of its ATPase activity by ESCRT-III proteins.

To assess whether spastin can extract CHMP1B from CHMP1B-IST1^NTD^ assemblies on membranes, we performed a GUV assay in which we incubated LD555-CHMP1B with IST1^NTD^ on GUVs containing 60 mol% egg-PC and 40 mol% DOPS for 30 min and then added it with spastin in solution (Fig. 4a). LD655-spastin bound to ESCRT-coated GUVs, but we did not observe any loss of fluorescence from CHMP1B, even after a 30 min of incubation in the presence of 1 mM ATP, suggesting that the ESCRT coat is intact and stably bound on the membrane, just as in the buffer control (Fig. 4b). Altogether our data suggests that, although recruitment to CHMP1B is robust, spastin did not uncoat CHMP1B from the membrane *in vitro*.

**Fig.4:**
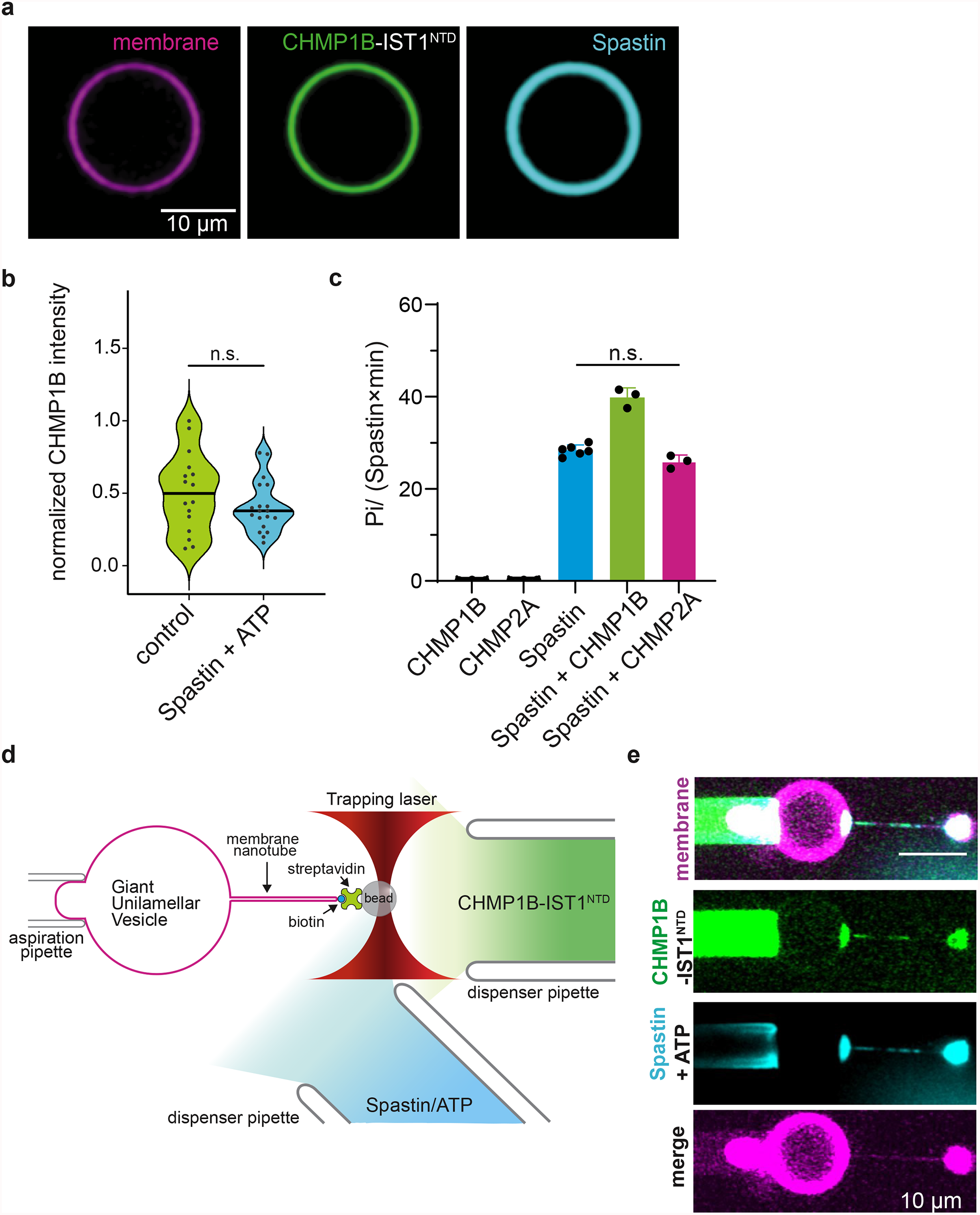
Spastin colocalizes with ESCRT-III enriched sites but does not uncoat nor sever the membrane. **a**, Representative image of GUVs labeled with DOPE-ATTO488 and pre-adsorbed with 500 nM LD555-CHMP1B and IST1^NTD^ after 30 min incubation at room temperature with 500 nM LD655-Spastin and 1 mM ATP. Scale bar is 10 μm. **b**, Violin plot of the distribution of LD555-CHMP1B fluorescence on the GUV after +/- 500 nM LD655-Spastin and 1 mM ATP. **c**, Spastin ATPase activity in the presence of ESCRT-III subunits. 2 μM ESCRT-III subunit were incubated with 0.2 μM spastin and 2 mM ATP for 10 min at 37°C. Spastin activity is mildly stimulated by CHMP1B but not with CHMP2A. At least three biological replicates were performed for each experimental condition. **d**, Schematic representation of the tube pulling assay geometry as previously described but with the addition of a third pipette to dispense spastin and ATP. **e**, Representative confocal images of LD655-spastin (cyan) colocalizing on LD555-CHMP1B (green) and IST1^NTD^ enriched sites on the membrane (magenta). 5 μM LD555-CHMP1B and 5 μM IST1^NTD^ were dispensed using a micropipette in proximity to the region of interest. 5 μM Spastin with 1 mM ATP was added after LD555-CHMP1B fluorescence equilibrated. No scission was observed in all of our trials (*n*=4). Scale bar is 10 μm.

### Spastin binding and ATPase stimulation by CHMP1B does not sever membrane nanotubes

We found that spastin hydrolyses ATP at a rate of 28 ± 1 ATP/spastin·min and this rate was slightly enhanced by soluble CHMP1B (Fig. 3c). There was no activation by a similar ESCRT-III protein, CHMP2A, which also has a VPS4-binding MIM domain (21, 22). Maximal ATPase activation of spastin was only observed in the presence of microtubules, consistent with its established function in microtubule severing (Extended Data Fig. 2). We next examined if this ATPase activity was required for scission of membranes bound with CHMP1B and IST1^NTD^ *in vitro* using the nanotube pulling assay. We introduced 5 μM spastin with 1 mM ATP using a third pipette at an angle close to the experimental region (Fig. 4d). This allows a controlled sequential delivery of spastin after initial binding of ESCRT-III proteins on preformed membrane nanotubes (Supplementary Video 3). Spastin bound immediately to ESCRT-III on the tube after delivery as evidenced by strong colocalization of LD655-spastin to CHMP1B on the tube in all of our trials (*n* = 4) (Fig. 4e). This observation is consistent with previous reports of spastin colocalization to CHMP1B enriched sites such as at the midbodies of dividing cells (51). However, in these experiments, carried out at constant tube length, we did not observe membrane scission even after more than 5 min (Supplementary Video 3). These data show that direct interaction of spastin and CHMP1B in the presence of ATP does not sever membranes.

### VPS4 uncoats membrane bound ESCRT-III proteins

Because VPS4 is the main ATPase tightly linked in canonical ESCRT processes we asked whether VPS4 could promote disassembly of CHMP1B-IST1^NTD^ analogous to VPS4 activity on CHMP2-CHMP3 filaments (38, 42, 54). VPS4 has two paralogs, VPS4A and VPS4B, in mammalian cells, which have essentially equivalent functions. We focused on the VPS4B isoform for all our studies since we found that it is easier to express and purify. Human VPS4B has undetectable ATPase activity in the absence of substrate proteins (Fig. 5a). We observed that VPS4B activity was only marginally increased by CHMP1B (3.4 ± 0.9 ATP/VPS4B·min), while incubation with CHMP2A produced ATP hydrolysis rate of 12.7 ± 0.4 ATP/VPS4B·min (Fig. 5a). Our data were consistent with previous findings that VPS4 engages the MIM domain of ESCRT-III proteins to stimulate its ATPase activity (21, 22, 55, 56).

**Fig.5:**
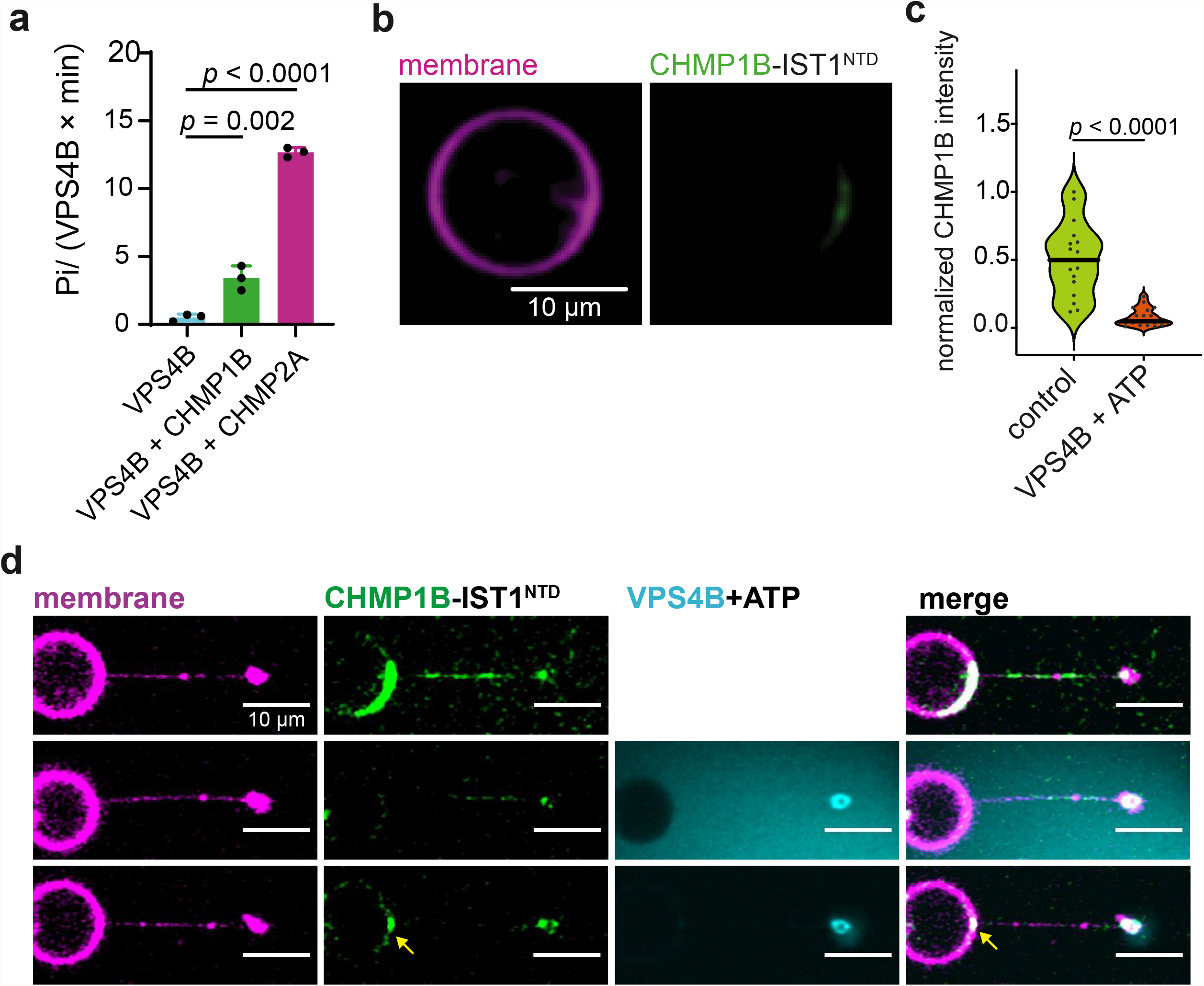
VPS4 uncoats ESCRT-III but do not sever the membrane. **a**, Rate of ATP hydrolysis by human VPS4B in the presence of full-length CHMP1B and CHMP2A. 2 μM ESCRT-III subunit were incubated with 0.2 μM spastin and 2 mM ATP for 10 min at 37°C. At least three biological replicates were performed for each experimental condition. **b**, Representative image of GUVs labeled with DOPE-ATTO488 and pre-adsorbed with 500 nM LD555-CHMP1B and IST1^NTD^ after 30 min incubation at room temperature with 500 nM VPS4B and 1 mM ATP. Scale bar is 10 μm. **c**, Violin plot of the distribution of LD555-CHMP1B fluorescence on the GUV after +/- 500 nM VPS4B and 1 mM ATP. **d**, Snapshots of LD655-VPS4B and ATP disassembling LD555-CHMP1B and IST1^NTD^ from the membrane (magenta) without severing the tube. 5 μM LD555-CHMP1B and 5 μM IST1^NTD^ were dispensed using a micropipette in proximity to the region of interest. 5 μM VPS4B with 1 mM ATP was added after LD555-CHMP1B fluorescence equilibrated. No scission was observed in all of our trials (*n*=3). Scale bar is 10 μm.

We sought to determine if VPS4 could disassemble and unfold CHMP1B-IST. Treatment of pre-adsorbed LD555-CHMP1B and IST1 on GUVs with VPS4B and ATP led to uncoating of ESCRT proteins from the membrane, in contrast to GUVs with VPS4B omitted (Fig. 5b, 5c). These findings confirm that VPS4B engages MIM1 containing ESCRT-III proteins and disassembles them, as expected.

### ESCRT disassembly by VPS4 does not lead to membrane nanotube scission

VPS4 is required for reverse topology scission by ESCRTs (16). We asked the presence of VPS4 and ATP could lead to scission using the tube pulling assay. To test this, we delivered 5 μM VPS4B and 1 mM ATP simultaneously at a low flow rate using a micropipette to pre-pulled nanotubes from GUVs after initial CHMP1B-IST1^NTD^ binding. Upon addition of VPS4B and ATP, we observed immediate loss of LD555-CHMP1B fluorescence intensity from the membrane, suggesting that the ESCRT protein was being uncoated (Fig. 5d). However, we observed rearrangement of LD555-CHMP1B intensity accumulating as a puncta at the junction of the tube and GUV (Fig. 5d) that seems refractory to VPS4/ATP treatment. This could be due to aggregation of CHMP1B and IST1^NTD^ that might hinder access by VPS4B. We did not see any evidence of membrane scission by VPS4B uncoating of CHMP1B from the membrane in any trial. These data suggests that VPS4 is not necessary for membrane scission on normal topology scission by CHMP1B and IST1 *in vitro*, although it is involved in recycling of ESCRT-III back to the cytosolic pool.

## Discussion

The membrane nanotube reconstitution reported here complements the recent cryo-EM study of Nguyen et al. (24), which demonstrated that addition of IST1 could strongly constrict CHMP1B tubes without leading to scission. By establishing conditions under which these constricted tubes can be severed, our observations provide a holistic account of the action of ESCRT-III proteins CHMP1B and IST1 in the constriction and scission of positively curved membrane tubules. Both the cryo-EM and membrane nanotube studies find that the CHMP1B-IST1 coat strongly constricts membrane tubes but the constriction, by itself, does not lead to scission. The degree of constriction observed differed between the two studies. Here, we did not observe tube radii below 15 nm and therefore not sufficient to provide elastic energy to drive membrane scission (57), consistent with the absence of scission in the static tube coating experiments. Our experiments used naturally occurring phosphatidylcholine with a mixture of 16:0, 18:0, 18:1, and 18:2 tails. Using cryo-EM, Nguyen et al. (24) reported that the gap between the inner leaflets of membrane tubes was narrowed by CHMP1B-IST1 coating to ∼5 nm. In order to achieve the highly constricted protein-coated tubules in the cryo-EM study, the asymmetric polyunsaturated (18:0 and 22:6) lipid, SDPC (58, 59) was used. Using brominated labels, this team found that the ends of the SDPC tails could be found near the membrane surface (60). The exceptional malleability of SDPC (59) probably explains why such a high degree of constriction could be observed in the cryo-EM study. Our study shows that CHMP1B-IST1 coat is also capable of constricting membrane tubes of a more physiological composition, although it constricts them to a lesser degree.

We were also able to answer a long-standing question concerning the role of VPS4 and spastin in membrane scission by CHMP1B and IST1. We found that VPS4 disassembles the CHMP1B-IST1 coat on the tubes, as expected. The major new finding here is that VPS4 does not lead to scission in this reaction, nor is it required for friction driven scission. The main role of VPS4 in reverse topology scission is thought to be recycling of the CHMP2 cap to allow further rounds of CHMP4 polymerization (42). Since our system does not contain CHMP2, it is not surprising that VPS4 is not required for CHMP1B and IST1 polymerization on positively curved membranes. We asked whether spastin, like VPS4, could recycle CHMP1B subunits. The primary function of spastin in the cell is to sever microtubules in an ATP-dependent manner (44, 61-63). The recruitment of spastin via MIT interaction with MIM-containing ESCRT-III subunits and its ATPase activity are both essential for the regulation of endosomal tubules (25-27). We asked if spastin could have a second role in disassembly of CHMP1B and IST1 from the membrane (52, 53). Our finding that spastin localizes on CHMP1B and IST1 coated tubes was expected given the tight interaction of spastin with CHMP1B. However, this interaction does not disassemble the coat nor sever the membrane. These data support the “standard model” that spastin is a microtubule severing enzyme that can be localized within cells by ESCRTs (27, 51, 64), but is not itself an ESCRT disassemblase.

We found that membrane scission occurs on CHMP1B-IST1 coated tubes when an additional mechanical pulling force is applied. Pulling velocities of the order of 3 μm s^-1^ were used in this study, which exceed the minimum value of 1 μm s^-1^ previously reported for FDS by the endophilin coat (49). Pulling of the membrane under a rigid protein scaffold in this velocity regime leads to friction between the lipids and the scaffold. Exposed hydrophobic amino acid side chains from helix α1 of CHMP1B were found by molecular simulations based on the cryo-EM structure (24) to lead to “scoring” of the membrane surface and exposure of phospholipid hydrocarbon tails (60). These strong interactions presumably account for the friction between the inner surface of the CHMP1B coat and the membrane. A friction driven scission (FDS) mechanism has been proposed to explain dynamin-independent vesicle release in by the N-BAR domain protein endophilin (49, 65). In FDS, molecular motors operating on microtubules, such as dynein, contribute the pulling force. Scission occurs when membrane tubules are destabilized by micropores created when a protein scaffold prevents the equilibration of membrane tension by limiting the lipid flow underneath upon tube extension (49). Microtubules are closely connected to the biological pathways that involve CHMP1B-IST1 coating of tubular vesicles (25-29). Thus, it seems reasonable to hypothesize that such a process could account for membrane scission in these cases.

Our findings lead us to propose the following model for the role of spastin and VPS4 in normal topology membrane scission by ESCRT-III proteins (Fig. 6). Recruitment of CHMP1B to the membrane provides initial constriction and IST1 binding further drives this constriction (Fig. 6a, 6b). Simultaneously or subsequently, addition of some external force applied by microtubule-associated motor proteins, which lead to a frictional force and membrane scission (Fig. 6c). In parallel, recruitment of spastin to ESCRT-III enriched sites severs the microtubules that run parallel with endosomal recycling tubes (51) (Fig. 6d); meanwhile, VPS4 uncoats CHMP1B and IST1 from the membrane back to the cytoplasmic pool (Fig. 6e). It remains to be seen what prevents spastin microtubule severing activity and VPS4 mediated ESCRT disassembly until the necessary FDS reaction has been completed. This could be under kinetic control, or there could be additional levels of regulation to be discovered. These insights establish a second instance of FDS, previously reported only for endophilin (49), and establish a biophysical basis for normal topology membrane severing by ESCRTs.

**Fig.6:**
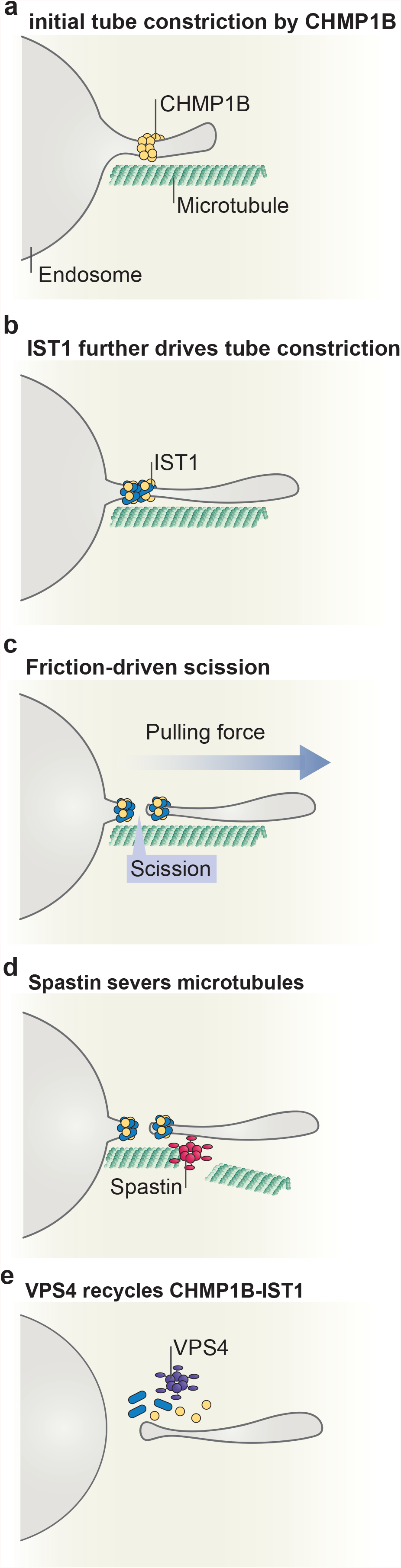
Model for friction-driven scission by CHMP1B and IST1. **a**, CHMP1B binds on the outer leaflet of endosome tubules forming a scaffold that constricts the positively-curved tube. **b**, As the tube continues to grow, IST1 gets recruited to the membrane forming an assembly with CHMP1B that further constricts the tube. **c**, Tube elongation promoted by an external pulling force can induce scission by friction between the protein scaffold and the underlying membrane. **d**, Spastin gets recruited to ESCRT-III enriched sites and severs the microtubules surrounding the tube. **e**, Finally, the AAA+ ATPase VPS4 disassembles the ESCRT-III assembly back to the cytosol.

## Materials and Methods

### Reagents

The following lipids: L-α-phospatidylcholine (EPC, 840051C), 1,2-dioleoyl-*sn*-glycero-3-phosphocholine (DOPC,850375C), 1,2-dioleoyl-*sn*-glycero-3-phospho-L-serine (DOPS, 840035C), 1,2-distearoyl-*sn*-glycero-3-phosphoethanolamine-N-[biotinyl(polyethylene glycol)-2000] (DSPE-PEG(2000)Biotin, 880129C) were purchased from Avanti Polar Lipids (Alabaster, AL) while 1,2-dioleoyl-*sn*-glycero-3-phosphoethanolamine-ATTO488 (DOPE-ATTO488, AD488-161) was purchased from ATTO-TEC. Lipid stock concentrations of 5 mg/mL were prepared by addition of chloroform from initial stocks of 10 mg/mL except for DOPE-ATTO488 where the lipid were solubilized in chloroform to a final concentration of 1 mg/mL. All lipids were kept under argon to avoid lipid oxidation and stored in -20°C in amber vials.

Streptavidin-coated silica beads (diameter 1.56 μm) were purchased from Spherotech (Lake Forest, IL). All other reagents were purchased from Sigma-Aldrich (St. Louis, MO, USA) unless otherwise specified.

### Protein expression, purification, and labeling

A single cysteine mutation at position 2 was introduced in the full-length human CHMP1B (CHMP1B^S2C^), for fluorophore labeling with maleimide dye, and IST1 containing residues 1-189 (IST1^NTD^) were fused to a TEV cleavable N-terminal His_6_-MBP tag. CHMP1B^S2C^ and IST1^NTD^ were transformed into *Escherichia coli* strain Rosetta (DE3) pLysS and grown in LB medium at 37°C. Upon reaching 0.6 at OD_600_ cells were induced with 0.5 mM isopropyl β-D-1-thiogalactopyranoside (IPTG) for 16 h at 20°C. Cells were pelleted, resuspended, and lysed in lysis buffer (50 mM Tris-HCl, pH 8.0; 500 mM NaCl; 20 mM imidazole; 0.5 mM tris(2-carboxyethyl)phosphine (TCEP) supplemented with 1 mM phenylmethylsulfonyl fluoride (PMSF) and EDTA-free protease inhibitors (Roche). The lysate was clarified by centrifugation and the proteins in the supernatant were purified by gravity Ni-nitrilotriacetic acid (Ni-NTA) affinity chromatography. The bound proteins were washed extensively with lysis buffer (lacking protease inhibitor) and eluted with elution buffer (50 mM Tris-HCl, pH 8.0; 500 mM NaCl; 250 mM imidazole; 0.5 mM TCEP). The eluate was concentrated and then applied to an initial size-exclusion chromatography (SEC) step using HiLoad 16/60 Superdex 200 (GE Healthcare, Chicago IL) in SEC200 buffer (50 mM Tris-HCl, pH 8.0; 150 mM NaCl, and 0.5 mM TCEP). Peaks corresponding to target protein were pooled and diluted 1.2-1.5 fold with SEC buffer followed by TEV cleavage at room temperature for 2 h. The pooled fractions were passed through a fresh 5 mL Ni-NTA resin to capture the His_6_-MBP tag, TEV, and uncleaved proteins. The eluate was concentrated and applied to an SEC step using HiLoad 16/60 Superdex 75 (GE Healthcare, Chicago IL) in SEC75 buffer (50 mM Tris-HCl, pH 7.4; 150 mM NaCl, and 0.5 mM TCEP). Highest peak corresponding to the target protein were pooled and concentrated. 2x molar excess LD555 or LD655 (Lumidyne Technologies) were added to fluorescently label the protein. The protein + dye solution was rotated gently in a nutator at room temperature for 2 h and then at 4°C overnight. Free dye were removed by passing the labeled protein through two 10DG desalting columns (Cytiva) and then through a Superdex75 10/300 size exclusion column. Proteins were flash frozen in liquid nitrogen and stored in -80°C freezer until use.

Codon-optimized synthetic gene for M87 Spastin with a single engineered cysteine at position 2 (^M87^Spastin^A2C^) were fused to an N-terminal TEV cleavable His_6_-GST tag and transformed and expressed in *Escherichia coli* strain Rosetta (DE3) pLysS. Expression condition was similar as described above. Cells were lysed by sonication in GST lysis buffer (50 mM Tris-HCl, pH 8.0, 500 mM NaCl, 5 mM MgCl_2_, 0.5 mM TCEP, 10% glycerol, and 1 mM ATP) supplemented with 1 mM PMSF and EDTA-free protease inhibitor cocktail. The cleared lysate was applied into glutathione-Sepharose 4B resin (GE Healthcare, Piscataway, NJ) overnight in 4°C on a nutator. The resin was washed extensively with GST lysis buffer followed by incubation with 1 mM ATP to release contaminating bacterial ATPases, and then eluted with GST lysis buffer supplemented with 25 mM reduced glutathione. The eluent was diluted with IE-A buffer (50 mM Tris-HCl, pH 8.0, 0.5 mM TCEP) to have a final NaCl concentration of 75 mM before being applied to a 5 mL HiTrap SP HP cation chromatography column (Cytiva, Marlborough, MA). The protein was eluted with a linear gradient of 75 mM to 1 M NaCl (IE-B, 50 mM Tris-HCl, pH 8.0, 1M NaCl, 0.5 mM TCEP). TEV was added to the pooled fractions and cleaved for 3 h at room temperature and passed through 5 mL Ni-NTA resin to capture TEV, His_6_-GST tag, and uncleaved protein. Eluate was concentrated and applied to SEC HiLoad 16/60 Superdex 75 column. Highest peak corresponding to ^M87^Spastin^A2C^ were pooled, concentrated, flash frozen in liquid nitrogen, and stored in -80°C freezer until use. Protein labeling with LD655 were performed as described above.

Human VPS4B with single mutation at position 2 for maleimide fluorophore labeling (VPS4B^S2C^) were transformed and expressed as described above. Cells were lysed by sonication using lysis buffer with protease inhibitor cocktail (50 mM Tris-HCl, pH 8.0, 500 mM NaCl, 5 mM MgCl_2_, 20 mM imidazole, 0.5 mM TCEP, 10% glycerol, and 1 mM ATP) and applied into Ni-NTA beads. The bound protein on the resin was washed excessively with lysis buffer and incubated with 1 mM ATP for 30 min at room temperature to release contaminating bacterial ATPases. The protein was eluted with lysis buffer containing 250 mM imidazole. The NaCl concentration of the eluate was adjusted to 75 mM with IE-A buffer and applied to a 5mL HiTrap Q Sepharose anion exchange column (GE Healthcare) and eluted with a linear gradient of 75 mM to 1 M NaCl (IE-B buffer). Fractions corresponding to VPS4B^S2C^ were pooled and labeled with 2x molar excess LD655 overnight. Excess dye was removed as described above. Protein was further purified by SEC using a HiLoad 16/60 Superdex 75 column. Protein was concentrated, flash frozen, and stored as described previously.

### Giant unilamellar vesicle (GUV) preparation

GUVs were prepared by hydrogel-assisted swelling as previously described. Briefly, 25 × 25 #2 coverslips were thoroughly rinsed while sonicating in water then ethanol then methanol and finally in water. 150 μL of 5% (w/v) polyvinyl-alcohol with molecular weight of 145,000 (Millipore) were spin-coated on plasma-cleaned coverslips. The thin polymer film was dried in an oven set to 60°C for 30 min. 15 μL of a 1 mg/mL lipid solution with the following mixture: 59.4% ePC, 40% DOPS, 0.5% DOPE-ATTO488, 0.1% DSPE-PEG-Biotin were spread uniformly on the slide using a Hamilton syringe. The lipid film was dried for 1 h under high vacuum to remove excess solvent and rehydrated with growth buffer (20 mM Tris-HCl, pH 7.4; 40 mM NaCl; 160 mM sucrose) at room temperature for 45 min – 1 h. GUVs were harvested by pipetting from the slides and were used immediately.

### Optical tweezers and micropipette manipulation integrated to a confocal microscope

We made modifications from a previous custom-built optical trap on an inverted Nikon Ti-Eclipse microscope used to perform all our membrane tube pulling assays (16). Briefly, we use a collimated 5 W 1070 nm continuous-wave infrared laser beam (YLR-5-1080-Y12, IPG Photonics) tightly focused on the image plane by a Plan Apochromat 60X 1.2 numeric aperture (N.A.) water immersion objective (Nikon). The laser power of the trap was controlled by a half-wave plate introduced in the optical train followed by a polarizing beam splitter (PBS) cube prior to entering the back aperture of the objective. Before it enters the confocal microscope body the beam is expanded with lenses in a 4f-configuration to slightly overfill the back aperture of the objective. The beam is then guided to the back port of the confocal microscope via two mirrors forming a periscope. The position of the trap in the z-axis was adjusted to match the z-focus of the scanning confocal unit. The position of the bead relative to the center of the trap was monitored using back focal plane interferometry (BFP), by imaging the BFP of the condenser on a quadrant photodiode (QPD) system (E4100, Elliot Scientific). The instrument was controlled and raw voltage data from the QPD were converted to trapping forces in real time using custom software written in LabVIEW. The trap stiffness (0.1-0.2 pN/nm range) was obtained from the Lorentzian fit to the power spectrum of position fluctuations of the trapped bead. Bright field illumination was provided by the top condenser unit and wide field fluorescence imaging light was introduced from the back port of the microscope (Sola SE II 365, Lumencor) to a video camera positioned to the right camera port. Fluorescence imaging was controlled by NIS-Elements imaging software (Nikon).

Micromanipulators (MPC-145, Sutter Instrument) were attached on each side of the microscope stage and were used to hold micropipettes. A custom open chamber holder was machined to accommodate insertion of micropipettes on either side of the chamber. Pressure inside the micropipettes was controlled by a high precision pressure controller (MFCS-EZ, Fluigent).

### Malachite green ATPase activity assay

All enzyme activity were determined by measuring the release of phosphate and formation of a green complex formed between malachite green oxalate with phosphomolybdate and using the malachite green assay as previously described (55) with slight modifications and manufacturer specifications (Sigma-Aldrich). 40 μL of ^M87^Spastin^A2C^ and VPS4B^S2C^ in ATPase buffer (20 mM Tris-HCl, pH 7.4, 100 mM KCl, 5 mM MgCl2, 0.5 mM TCEP) were aliquoted into a 96-well plate. Addition of 40 μL ESCRT-III protein or microtubule substrate in ATPase buffer supplemented with or without 2 mM ATP initiates the reaction. Final enzyme and protein concentrations in the reaction were 0.2 μM and 2 μM respectively. The reaction mixture was incubated at 37°C for 10 min and terminated by addition of 20 μL malachite green reagent. The reaction was further incubated for 60 min at room temperature prior to measuring absorbance with a plate-reading luminometer (GloMax, Promega). Samples were correlated to phosphate concentration standard controls. At least three biological replicates were performed for each experimental condition.

### Fluorescence ATPase activity assay of ESCRTs on the GUV

GUVs were incubated with LD555-CHMP1B^S2C^ and IST1^NTD^ at a final concentration of 500 nM in experimental buffer (20 mM Tris-HCl, pH 7.4; 100 mM NaCl, 40 mM glucose, and 0.5 mM TCEP) for 15 min at room temperature. An eight-well observation chamber (Lab-Tek) was passivated with 5 mg/mL β-casein dissolved in experimental buffer for 30 min and rinsed three times. GUVs were then mixed with either spastin or VPS4B (final [enzyme] = 500 nM) in experimental buffer supplemented with 1 mM ATP and 1 mM MgCl_2_ at room temperature in the observation chamber. Reaction mixtures were imaged after 30 min. At least three biological replicates were performed.

### Fluorescence recovery after photobleaching (FRAP) of ESCRTs and membranes

GUVs with pre-bound ESCRTs were prepared similar to the above description and were subsequently diluted 5X in experimental buffer to avoid recovery from soluble protein in the GUV external solution. FRAP was performed with the excitation λ = 561 nm laser for LD555-CHMP1B^S2C^ or λ = 650 nm laser for DOPE-ATTO647 labeled membrane on a Nikon A1 confocal microscope with a 60X Plan Apochromat 1.4 N.A. oil immersion objective. A small rectangular region on the GUV with pre-bound ESCRTs was imaged at low laser power (30 μW) for 3-5 sec followed by 100% laser (15 mW) power for 15-30 seconds. Fluorescence recovery was imaged at low laser power over the course of 3 min. At least three biological replicates were performed.

### Micropipette preparation

Micropipettes made of borosilicate capillaries (ID =0.78 mm, OD = 1 mm, B100-75-15, Sutter Instrument) were formed using a puller (P-1000, Sutter Instrument) and the tips were then microforged (MFG-5, MicroData Instrument, NJ USA) to an internal diameter of 4-5 μm for the GUV holder and 7-10 μm for dispenser pipettes.

### Membrane nanotube pulling assay

An open experimental chamber was formed by using two clean glass coverslips (VWR) separated by 1 mm on a custom-built chamber holder. The coverslips and interior of the micropipette used to hold the GUV were passivated with 5 mg/mL β-casein dissolved in experimental buffer for 30 min and rinsed three times and filled with experimental buffer. A micropipette was filled with fluorescently labeled protein using a MicroFil flexible needle (MF34G-5, World Precision Instruments, UK). The micropipettes were introduced in each side of the chamber and placed in the middle using micromanipulators. A third micropipette containing either spastin or VPS4B +/- ATP was inserted at an angle and all three pipettes were aligned in the same field of view. The pressure inside the pipette was adjusted so that no positive or negative flow is experienced inside the chamber. GUVs and streptavidin-coated silica beads were deposited in the chamber and allowed to settle. To form a membrane nanotube, a bead was held in place by an optical trap and put in contact with a GUV aspirated on the pipette and held at low tension and then subsequently pulled away. Formation of the membrane nanotube was verified by fluorescence between the GUV and the bead and the sudden increase in force as measured by the optical trap. The chamber was sealed on each side by adding mineral oil to avoid evaporation.

Once a tube is formed, a micropipette is lowered into the field of view and protein was allowed to flow gently without disturbing the tube. After protein binding is established, the micropipette is raised up and another micropipette containing either spastin or VPS4 is lowered and injected into the field of view. All proteins were adjusted to match the osmolarity of the experimental buffer using an osmometer (Osmette II, Precision Instruments).

### Cryo-electron microscopy (cryo-EM) sample preparation

Lipids to form large unilamellar vesicles (LUVs) consisting of (55% SDPC, 20% DOPS, 10% brain PI(4,5)P_2_, and 15% cholesterol, 2 mg total lipid) were dried in a round-bottom flask using a rotary evaporator at 55°C. The resulting thin lipid film was further dried inside a vacuum desiccator overnight. Liposomes were prepared by rehydrating the lipid with buffer (20 mM HEPES, pH 7.5 and 100 mM NaCl) to a final concentration of 2 mg/mL and sonicated for 1 h at 55°C followed by 10 freeze/thaw cycles. Lipids were then extruded 21 times through a 400 nm pore size filter, or until the lipid suspension begins to clarify, producing LUVs. LUVs (0.5 mg/mL) were mixed with 10 μM CHMP1B and IST1^NTD^ and incubated overnight at room temperature.

### Cryo-EM data collection

Incubated sample (3.5 μL) was applied to glow-discharged C-Flat Holey Carbon Grids (1.2/1.3, Au, 300 mesh). Grid blotting and vitrification were performed using a Vitrobot Mark IV (Thermo Fisher Scientific) (100% humidity, 22°C, blot time = 5s, blot force = 5) and Whatman 595 blotting paper.

Vitrified samples for cryo-EM were screened on a Talos Arctica (Thermo Fisher Scientific) with a Gatan K3 Summit direct detection camera in super-resolution counting mode with a pixel size of 0.4495 Å. Data were manually collected using SerialEM (66) and spanned a range of 2-5 μM defocus. Movies consisted of 50 frames, with a total dose of 85.2 e^−^/Å^2^ and a total exposure time of 10s. Movie frames were motion-corrected and dose-weighted using MotionCor2 (67).

## Acknowledgements

We would like to thank S. Shukla, K. Rose, and L. Jensen for critical reading of the manuscript.

## Extended Data Figure Legends

**Extended Data Fig.1:**
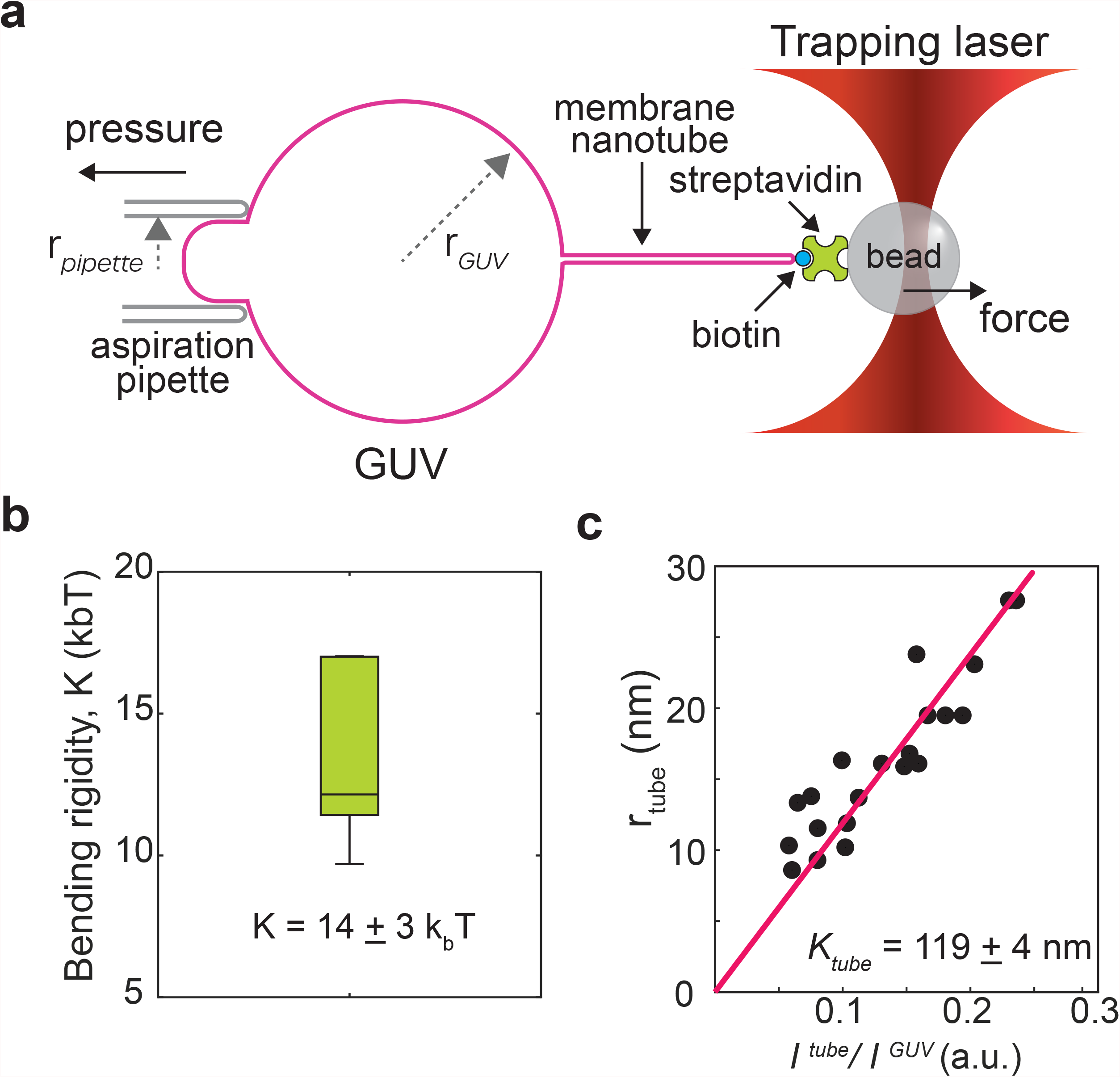
Tube radius measurement. **a**, Schematic of tube radius, in the absence of proteins, using an optical trap to extract force and membrane tension, σ, from the relationship 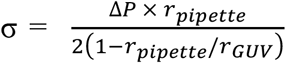, where rpipette is the pipette radius, rGUV is the GUV radius, and ΔP is the difference in aspiration pressure. **b**, Bending rigidity, K, is determined to be 14 ± 3 kbT by plotting the linear relationship between force and 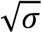. **c**, We determined tube radii measurements using the ratio between the fluorescence intensity of the GUV, I_tube_ and the fluorescence intensity of the membrane nanotube, I_GUV_ multiplied by a constant K_tube_ as described in (47). Ktube is experimentally determined by plotting the tube radius of a bare GUV (*N*=9) calculated from the expression rtube = F/4πσ, where σ is the membrane tension and F is the force measured using the optical trap and plotting against I_tube_/I_GUV_. We find the calibration constant Ktube = 119 ± 4 nm.

**Extended Data Fig.2:**
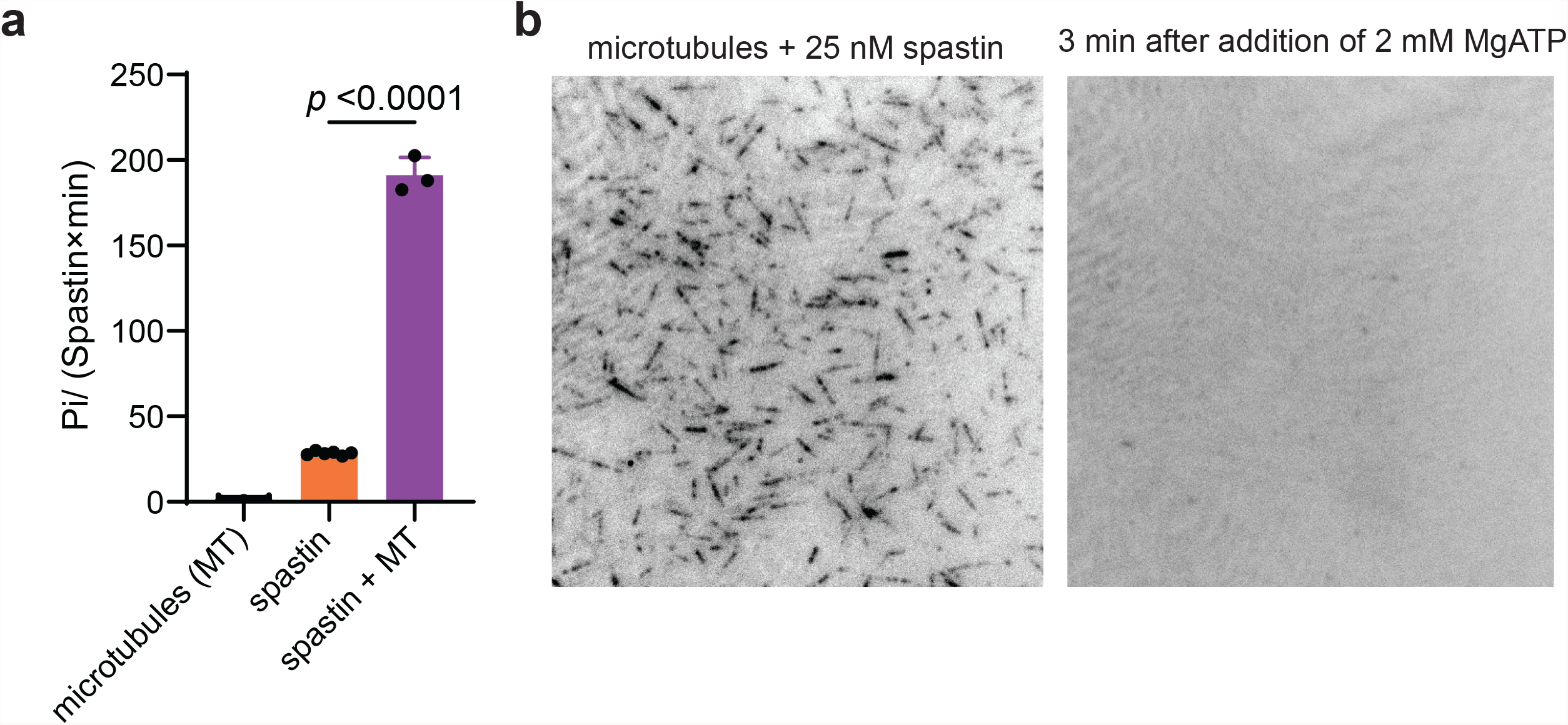
Spastin disassembles microtubules in an ATP-dependent manner. **a**, Spastin activity is stimulated in the presence of microtubules (191 ± 10 ATP/spastin·min). **b**, Fluorescently-labeled microtubules are immobilized on a coverglass together with spastin. Addition of ATP disassembles microtubules.

**Supplementary Video 1: Tube elongation in the absence of protein scaffold**. Bare membranes pulled at > 25μm.s^-1^ at 0.2 pN.nm^-1^ do not break even after repeatedly being brought back-and-forth.

**Supplementary Video 2: External pulling on protein-scaffolded tubes promotes scission** Membrane tube pulled at 3 μm.s^-1^ on membrane bound LD555-CHMP1B-IST1^NTD^ (green) protein scaffolds induces scission. Black arrow highlights the point of scission.

**Supplementary Video 3: Spastin colocalizes with ESCRT-III without uncoating or severing the tube**. 5μM LD555-CHMP1B and 5 μM IST1^NTD^ were dispensed using a micropipette in proximity to the region of interest. 5 μM Spastin with 1 mM ATP was added after LD555-CHMP1B fluorescence equilibrated. LD655-spastin (cyan) colocalizing on LD555-CHMP1B (green) and IST1^NTD^ enriched sites on the membrane (magenta).

